# Cellular and Viral Determinants of HSV-1 Entry and Intracellular Transport towards Nucleus of Infected Cells

**DOI:** 10.1101/2021.01.04.425350

**Authors:** Farhana Musarrat, Vladimir Chouljenko, Konstantin G. Kousoulas

## Abstract

HSV-1 employs cellular motor proteins and modulates kinase pathways to facilitate intracellular virion capsid transport. Previously, we and others have shown that the Akt inhibitor miltefosine inhibited virus entry. Herein, we show that the protein kinase C inhibitors staurosporine (STS) and gouml inhibited HSV-1 entry into Vero cells, and that miltefosine prevents HSV-1 capsid transport toward the nucleus. We have reported that the HSV-1 UL37 tegument protein interacts with the dynein motor complex during virus entry and virion egress, while others have shown that the UL37/UL36 protein complex binds dynein and kinesin causing a saltatory movement of capsids in neuronal axons. Co-immoprecipitation experiments confirmed previous findings from our laboratory that the UL37 protein interacted with the dynein intermediate chain (DIC) at early times post infection. This UL37-DIC interaction was concurrent with DIC phosphorylation in infected, but not mock-infected cells. Miltefosine inhibited dynein phosphorylation when added before, but not after virus entry. Inhibition of motor accessory protein dynactins (DCTN2, DCTN3), the adaptor proteins EB1 and the Bicaudal D homolog 2 (BICD2) expression using lentiviruses expressing specific shRNAs, inhibited intracellular transport of virion capsids toward the nucleus of human neuroblastoma (SK-N-SH) cells. Co-immunoprecipitation experiments revealed that the major capsid protein Vp5 interacted with dynactins (DCTN1/p150 and DCTN4/p62) and the end-binding protein (EB1) at early times post infection. These results show that Akt and kinase C are involved in virus entry and intracellular transport of virion capsids, but not in dynein activation via phosphorylation. Importantly, both the UL37 and Vp5 viral proteins are involved in dynein-dependent transport of virion capsids to the nuclei of infected cells.

**Importance:** Herpes simplex virus type-1 enter either via fusion at the plasma membranes or endocytosis depositing the virion capsids into the cytoplasm of infected cells. The viral capsids utilize the dynein motor complex to move toward the nuclei of infected cells using the microtubular network. This work shows that inhibitors of the Akt kinase and kinase C inhibit not only viral entry into cells but also virion capsid transport toward the nucleus. In addition, the work reveals that the virion protein ICP5 (VP5) interacts with the dynein cofactor dynactin, while the UL37 protein interacts with the dynein intermediate chain (DIC). Importantly, neither Akt nor Kinase C was found to be responsible for phosphorylation/activation of dynein indicating that other cellular or viral kinases may be involved.

## Introduction

### Cellular proteins involved in intracellular transport

Intracellular transport is mediated by the cytoskeleton network and motor proteins within the cytoplasm of cells. Short-distance movement of cargo is mediated by actin and myosin, while long-distance transport (>1μM) of organelles and other cargo requires ATP-dependent motor proteins and accessory factors, which are transported via the microtubule network (1–4). The plus end or the growing end of the microtubule is located towards the periphery (cell membrane) of the cell and helps in cell growth and migration. Several end-binding microtuble-associated proteins (MAPs) and plus-tip interacting proteins (TIPs) facilitate intracellular transport (5). Specifically, the EB1 protein recognizes the growing end of a dynamic microtubule and recruits other MAPs and facilitate binding of intracellular cargoes and organells to the microtubular network for intracellular transport (6–8). The minus end of the microtubule is proximal to the nucleus and the microtubule organizing center (MTOC). The two major motor proteins dynein and kinesin, which mediate intracellular cargo transport toward the nucleus and cell periphery, respectively. Dynein is a multi-subunit complex with each subunit having distinct functions. The heavy chains possess ATPase activity, which is required for movement along the microtubules. The intermediate chain (IC) and light chain (LC) regulate motor activity and mediate interactions with other cofactors and cargo (9–15). A number of different accessory proteins interact with the dynein motor complex and modulate dynein-dependent transport. Dynactin (DCTN) is a large molecule consisting of several components namely, DCTN1 (p150), DCTN2 (dynamitin/p50), DCTN3 (p24/22), DCTN4 (p62), DCTN5 (p25), DCTN6 (p27), CapZ and the Arp polymer. One end of the DCTN1 (p150) binds to the dynein IC and the other end is associated with microtubules. DCTN4 (P62), Arp polymer, and CapZ participate in cargo binding (16–19). The Bicaudal D homolog (BICD) facilitates dynein-dependent transport in epithelial and neuronal cells (20–22). BICD2 binds to DCTN2 (p50), docks on Arp polymer and helps in the recruitment of dynein cargo (23). BICD2 also interacts with the N-terminus of the dynein heavy chain and the light intermediate chain, which together enhance dynein-dynactin interactions (15). In addition, BICD2 is associated with the cellular transport of mRNA and organelles and helps to position the nucleus during cell division by docking at the nuclear pore (24).

### Viruses hijack dynein-dependent intracellular cargo transport to facilitate infection

Several viruses have been shown to utilize the cellular motor protein dynein to facilitate intracellular virion transport (25–30). HSV-1 capsids take advantage of cellular intracellular transport mechanisms to travel along cellular microtubules toward the cell nucleus where it delivers its genome for replication (7, 31–34). Although most of the outer tegument proteins of HSV-1 dissociate in the cytoplasm following entry into host cells, the tegument proteins UL36, UL37, and US3 remain attached to the capsid during intracellular transport to the nucleus and are known to be involved in intracellular capsid transport. (31, 33, 35–42). The HSV-1 inner tegument proteins UL36 and UL37 interact with the cellular motor proteins kinesin and dynein, respectively, Thes interactions have been suggested to cause bidirectional or saltatory movement along microtubules (31, 32, 36, 43, 44). In some cases, virus transport occurs at the expense of cellular transport. HSV-1 infection disrupts mitochondrial transport by altering kinesin-mediated motility (28). For pseudorabies virus (PRV), VP1/2 (UL36 homolog) interacts with the dynein intermediate chain (DIC) and dynactin (p150) in the HEK293 cell line, while VP1/2 alone was shown to move cargo via the microtubule network. The proline-rich sequence of VP1/2 appears to play an essential role in the retrograde transport of PRV in neuronal cultures and in vivo (45). Similar roles have been suggested for a phosphoprotein specified by pseudorabies and lyssavirus, which bind to the dynein light chain 8 (LC8). SiRNA-mediated depletion of the dynein heavy chain DYNC1H1 inhibited human immunodeficency virus (HIV) infection, although, siRNA depletion of other dynein components had little to no effect on HIV infection in TZM-bl cells (24). The dynein light chain 2 (DLC-2) transports HIV integrase and DLC-1 interacts with HIV capsid and affects reverse transcription (46). HIV-1 exploits one or more dynein adaptors for efficient infection and transport towards the nucleus (47). Dynein adaptor BICD2, dynactin components DCTN2, DCTN3 and ACTR1A (arp1 rod) were essential for HIV permissiveness in TZM-bl cells (24).The adenovirus hexon protein directly recruits the intermediate dynein chain (DIC) for capsid transport (48). Interestingly, dynein interaction with the simian virus 40 (SV40) virion after entry into cells causes a disassembly of the virion that is required for transport toward the nucleus (30). The tegument protein ppUL32 of human cytomegalovirus (HCMV) interacts with BICD1, and its depletion resulted in low virus yield due to insufficient trafficking of ppUL32 to the cytoplasmic viral assembly compartment (AC) (22). The end binding protein 1 (EB1) is essential for HSV-1 infection, and the microtubule plus end associated protein CLIP 170 initiates HSV-1 retrograde transport in primary human cells (7).

### Regulation of intracellular transport by kinases

Intracellular transport is regulated by various protein kinases through direct post-translational modification (phosphorylation) of motors, accessory proteins, cargoes, or via indirect modification of the microtubular network (49–57). Mitogen activated protein kinases (MAPKs) play a pivotal role in anterograde transport of neurofilament in differentiated neuroblastoma cells, and inhibition of MAPKs completely inhibits transport and organization of neurofilament in axonal neurites (58). The Akt (Protein kinase B) is involved in transport of synaptic vesicles, BDNF and APP containing secretory vesicles in cortical neurons (59). Protein kinase C (PKC) is involved in actin cytoskeleton modification in retinal bipolar cells and controls vesicular transport in primary human macrophages (54, 56). Also, PKC regulates intracellular organelle transport in sensory and sympathetic neurons. Activation of PKC and caspases by neurotoxin in dopaminergic neurons affect fast axonal transport, a phenomenon observed in Parkinson disease. In mice, PKC activation leads to inhibition of neurotrophin retrograde transport to the trigeminal ganglia and superior cervical ganglia when a PKC activator is injected in the eye (60–62). Some kinases directly modify motor proteins. Specifically, inhibition of glycogen synthase kinase (GSK3β) enhances retrograde motility by direct phosphorylation of dynein intermediate chain (DIC) (63). GSK3β also recruits EB1 to the plus end of microtubules, which is important for the loading of cargo and virions on the microtubular network (64).

### Modulation of cellular kinases involved in intracellular transport by viruses

Modification of cellular kinases is another way that viruses regulate intracellular transport. HSV-1 activates the serine-threonine kinase (Akt) (65) and mitogen activating protein kinases (MAPKs) at later times post entry (66). The PI3K/ Akt pathway is activated following HIV Env-CD4 mediated fusion. HIV requires PI3K p110α/PTEN signaling during entry into T cells, PM1 cells, and TZM-b1 cells (67). Inhibition of certain cellular kinases prevents HSV-1 entry and transport towards the nucleus. Specifically, Akt inhibition by miltefosine inhibits HSV-1 (McKrae) entry into Vero and SK-N-SH cells (68). Also, PI3K inhibition inhibits HSV containing endosome trafficking (69). Some viruses modify the motor proteins via activation of kinases for their transport in the cytoplasm. Adenovirus activates protein kinase A (PKA) and p38 MAPK and is involved in phosphorylation of the dynein light intermediate chain 1 (LIC-1, which subsequently binds the adenovirus hexon (70). Adenoviral activation of kinases enhances the nuclear targeting possibly through integrins (71).

In this report we investigated the role of cellular kinases and motor proteins in HSV-1 intracellular transport. Previously, we reported that Akt is necessary for HSV-1 entry into cells (72). Herein, we show that the broad spectrum Akt and PKC inhibitor miltefosine prevented HSV-1 entry and intracellular transport of HSV-1 capsids after virus entry into cells. We show that HSV-1 infection activates dynein by phosphorylating the intermediate chain (DIC) at early time post infection independently of PKC or Akt activation via phosphorylation. Furthermore, we show that the HSV-1 tegument protein UL37 interacts with the intermediate chain of cellular motor protein dynein (DIC) and that capsid protein VP5 (ICP5) interacts with motor accessory proteins dynactin (DCTN1/p150, DCTN4/p62), EB1 but not DIC during intracellular capsid transport towards the nucleus. These results suggest that HSV-1 modulates both virus entry and intracellular transport through specific protein-protein interactions and modulation of cellular kinases.

## Results

### Akt is required for HSV-1 (McKrae) intracellular capsid transport towards the nucleus

Previously, we showed that Akt inhibition blocke HSV-1 (McKrae) entry into epithelial and neuroblastoma cells (68). To determine if Akt has a role in intracellular transport of HSV-1, SK-N-SH cells were treated with the Akt inhibitor miltefosine at different concentrations (10μM, 30μM, 50μM, and 70μM) at 37°C and intracellular transport was monitored by immunofluorescence microscopy. Before drug treatment, cells were infected with HSV-1 (McKrae) (MOI 20) for 1 hour at 4°C and then shifted to 37°C for 15 minutes to allow virus entry into cells. Following a low pH buffer wash, cells were incubated with the inhibitors for 5 hours and subsequently, were processed for visualization of the ICP5 (Vp5) containing capsids by immunofluorescence. HSV-1 (McKrae) virions were transported toward the nucleus of infected cells resulting in productive infection, as evidenced by the expression of the ICP5 protein (red) inside the nucleus (blue) at 5 hpi. However, viral capsids remained in the periphery of the cytoplasm, in miltofesine-treated cells (Fig. 1A). At 15 minutes post infection, the relative locations of virion capsids detected were similar between “No Drug” and “Akt-treated cells” quantified by determining the average fluorescent signal per cell. In the presence of Akt, the fluorescence signals remained similar to the 15-minute time points for the Akt-treated cells (Fig. 1B).

**Figure 1.**
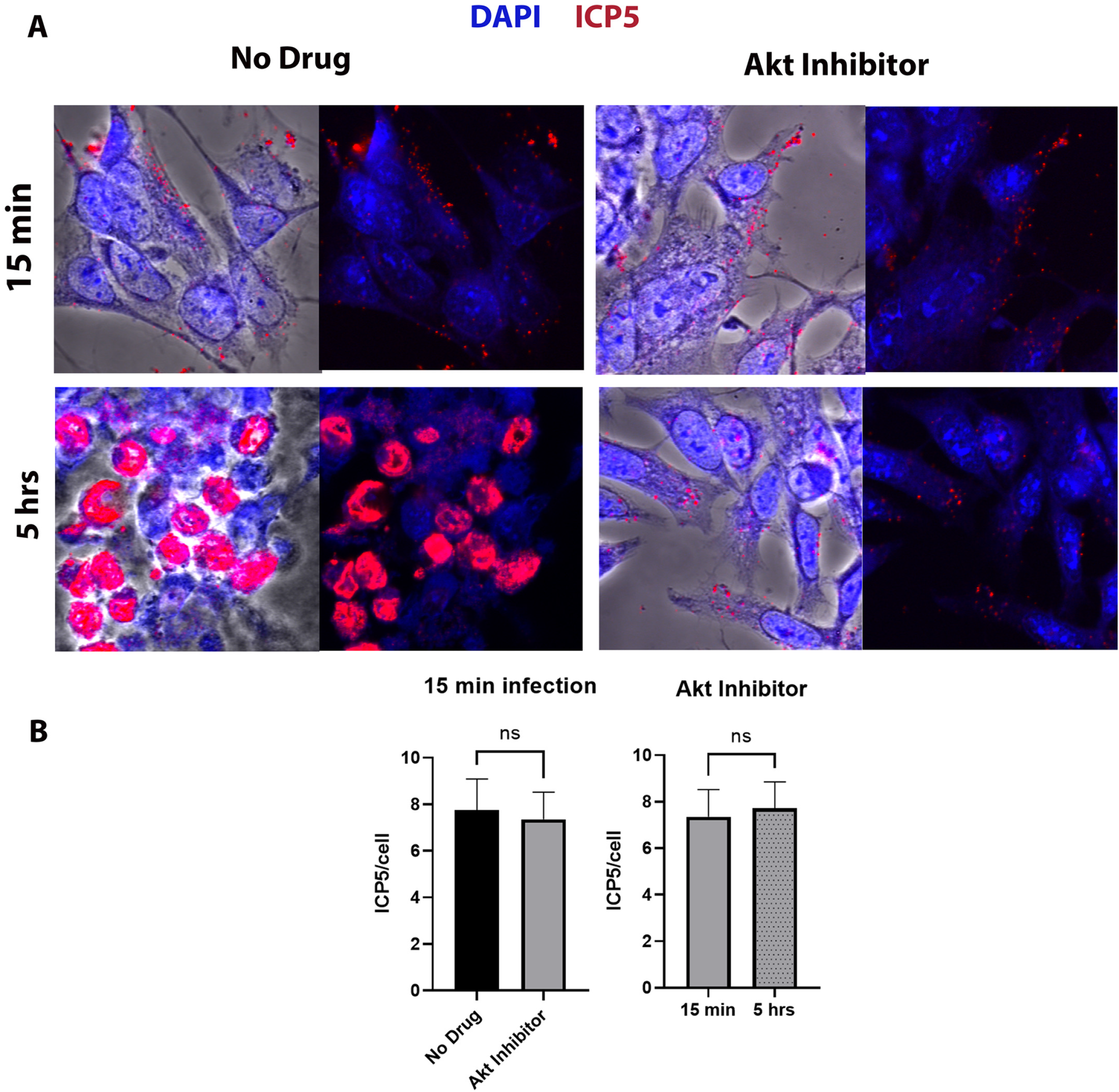
Effect of Akt HSV-1 transport. (A) SK-N-SH cells were adsorbed at 4°C with HSV-1 (McKrae) at MOI 20 for 1 hour then shifted to 37°C for 15 minutes, and then washed with low pH buffer. The Akt inhibitor miltefosine was added at a concentration of 30μM and incubated for 5 hours at 37°C. The 8 well chamber slide was then fixed with formalin, permeabilized with methanol, and prepared for fluorescent microscopy. Antibody against VP5 (ICP5) (red) was used, and nuclei were stained with DAPI (blue). Capsids colocalized in the nuclei appear purple. Magnification is 63X under oil immersion. (B) The amount of fluorescence detected with the anti-ICP5 antibody (virion capsids) was quantified by fluorescence imaging of 10 different random sections of slides, using ImageJ software. The fluorescence signals per cell were counted and their average value was used to plot the graph. Statistical comparison was conducted by Graph Pad prism using Students’ t-test. Bars represent 95% confidence interval about the mean. Differences were determined significant if P < 0.05.

### Protein kinase C (PKC) is required for HSV-1 (McKrae) entry

Our previous results indicated that the serine-threonine kinase Akt was required for HSV-1 entry into cells. Miltefosine blocks both Akt and PKC enzymatic activities. Therefore, we investigated the specific role of PKC in HSV-1 entry and transport in Vero cells. Staurosporine is a broad spectrum protein kinase inhibitor. It inhibits protein kinase C, protein kinase A, p60v-src tyrosine protein kinase and CaM kinase II at different concentrations. Gouml is a potent, fast acting and specific PKC inhibitor of different PKC subtypes such as PKCα, PKCβ, PKCγ, PKCδ, PKCζ and PKCμ. Treatment with either Gouml, or staurosporine for 4 hours at 37°C caused a dose-dependent reduction in the entry of HSV-1 (McKrae) (100 PFU) in Vero cells. Maximum inhibition was observed at 90μM (Gouml) and 0.5μM (STS) concentrations, respectively (Fig. 2A). Detection of the VP5 containing capsids via immunofluorescence revealed drastic reduction of capsids in the cytoplasm of infected cells (Fig. 2B). This reduction was the combined effect of these inhibitors in blocking both virus entry and intracellular capsid transport, since the incubation of infected cells for 4 hours at 37° C disloged virions bound to cell surfaces, but unable to enter (data not shown).

**Figure 2.**
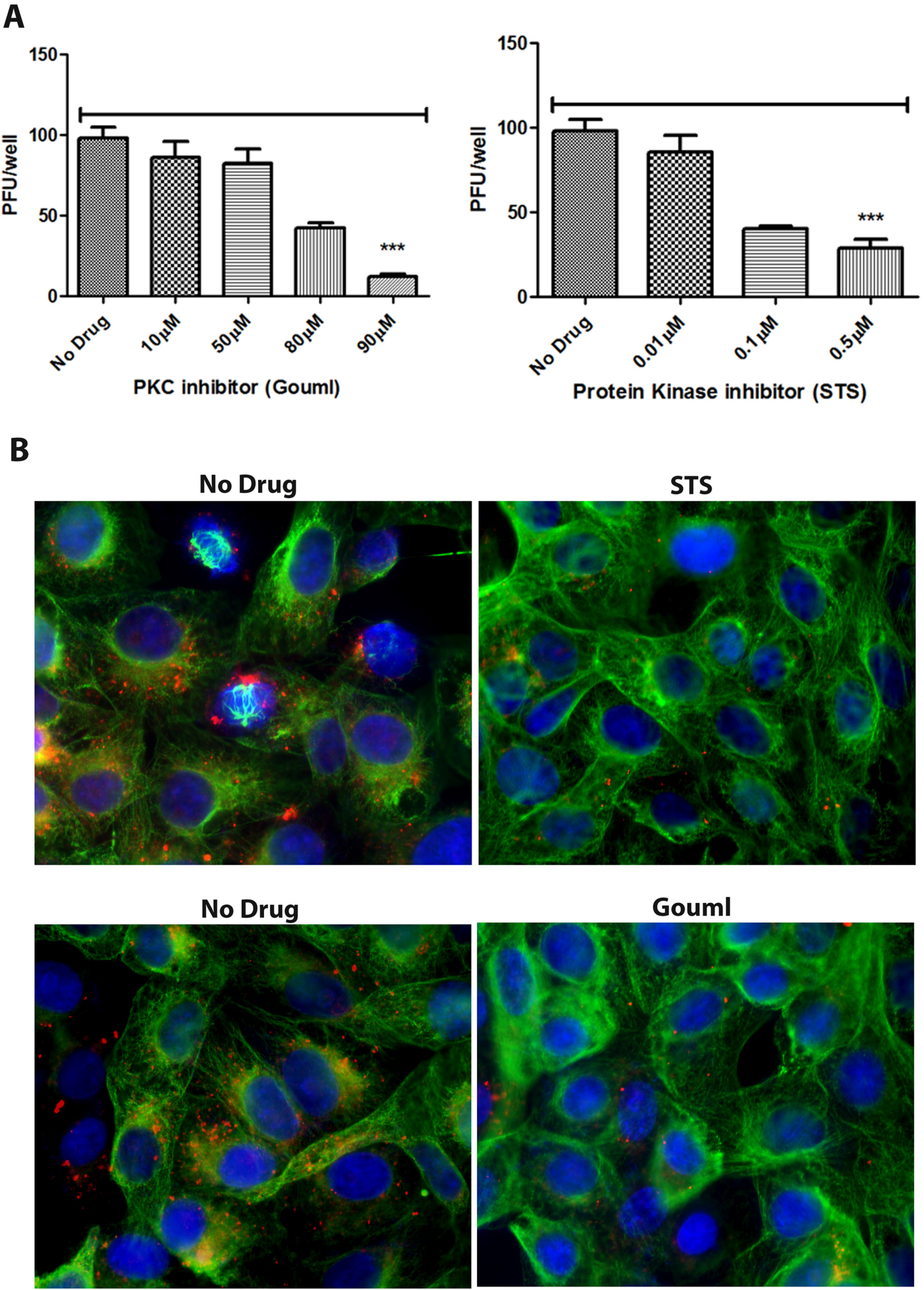
Effect of PKC inhibitors on HSV-1 entry. (A) Vero cells were treated with a series of dilutions of the PKC inhibitor (Gouml), or the protein kinase inhibitor (staurosporine STS) for 4 hours and then infected with HSV-1 (McKrae) (100 PFU) for 1 hour at 4°C. Cells were incubated at 37°C for 48 hours for plaque assay. Viral plaques were counted after crystal violet staining. ***P<0.001 vs. no drug-treated control. Statistical comparison was conducted by Graph Pad prism using ANOVA with post hoc t-test with Bonferroni adjustment. Bars represent 95% confidence interval about the mean. (B) Vero cells were treated with a series of dilutions of PKC inhibitor (Gouml) or the protein kinase inhibitor (staurosporine; STS) for 4 hours and then infected with McKrae (MOI 20) for 1 hour at 4°C. Cells were incubated at 37°C for 3 hours and then fixed with formalin and permeabilized with methanol. Antibodies against VP5 (ICP5) (red), and alpha-tubulin (green) were used, and nuclei were stained with DAPI (blue). Magnification 40X.

### HSV-1 (McKrae) activates the dynein intermediate chain (DIC)

Previously, it was shown that HSV-1 tegument proteins bind both plus and minus end directed motor proteins simultaneously, while moving along the microtubules (35, 36, 73). To determine if HSV-1 infection results in the modification of the dynein motor during intracellular transport, Vero cells were infected with HSV-1 (McKrae) and infected cell lysates prepared at different times post infection were analysed for dynein phosphorylation. HSV-1 infection caused phosphorylation of the intermediate dynein chain 1B as early as 15 minutes post-infection (Fig. 3A). An increase in the level of phosphorylation was observed from 15 minutes to 3 hours post-infection, which is approximately the time that it takes for HSV-1 capsids to be transported to the nucleus of infected cells (69).

**Figure 3.**
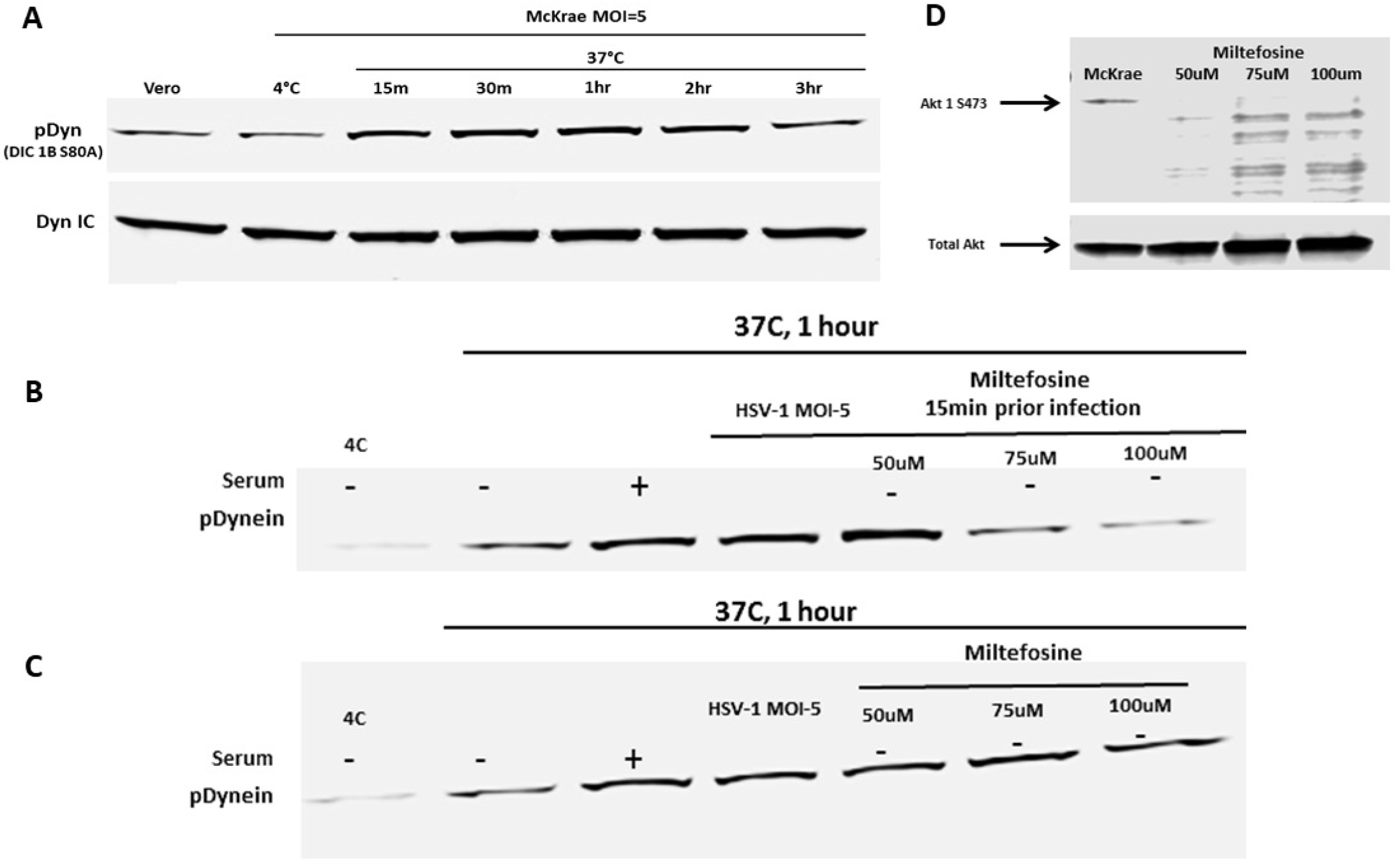
Dynein activation by HSV-1. (A) Serum-starved Vero cells were infected with HSV-1 (McKrae)at 4°C for 1 hour at MOI 5 and then shifted to 37°C for 15 m, 30 m, 1 hr, 2 hrs, and 3 hrs. Cells were lysed with the NP40 lysis buffer containing a protease inhibitor cocktail and analyzed for phosphorylated dynein intermediate chain (DIC 1BS80A) by SDS PAGE. The total dynein intermediate chain (DIC) protein was used as the loading control. (B) Vero cells were pre-treated with miltefosine at 37°C. Cells were infected with HSV-1 (McKrae) at 4°C for 1 hour at a MOI 5 and then shifted to 37°C for 1 hour. Cells were lysed with the NP40 lysis buffer containing a protease inhibitor cocktail and analyzed for the presence of the phosphorylated dynein intermediate chain (DIC 1BS80A) by SDS PAGE. (C) Following infection with HSV-1 (McKrae) at 4°C for 1 hour at MOI 5, cells were incubated at 37°C for 15 minutes, then miltefosine was added and incubated for 1 hour. Cell lysates were prepared the same way for SDS PAGE analysis. Serum + and – represents whether serum was added or not (D) Effect of miltefosine on Akt phosphorylation. Total Akt was used as loading control.

### Dynein phosphorylation/activation by HSV-1 (McKrae) is independent of either Akt or PKC signaling

Previously, we and others have reported that HSV-1 activated the Akt pathway at early times post-infection (65). We considered that the Akt/PKC pathway is upstream of dynein activation and plays a role in activating dynein for the capsid motility. To investigate if HSV-1 (McKrae) activates/phosphorylates dynein via the Akt/PKC pathway, miltefosine was used to treat cells before and after virus entry and the resultant dynein phosphorylation pattern was monitored at 1 hour post infection. Miltefosine inhibited dynein phosphorylation at the concentrations of 75 uM and 100 uM (Fig. 3B, 3D), which we reported to inhibit virus entry previously (72). However, once the virus entered the cell, blocking Akt or PKC did not inhibit activation of dynein by HSV-1 (McKrae) (Fig. 3C), but inhibited intracellular capsid transport (Fig. 1, Fig. 2B).

### Motor accessory proteins are essential for HSV-1 (McKrae) intracellular transport

To determine the role of motor proteins and adaptors in virion capsid intracellular transport towards the nucleus, we generated several SK-N-SH knockdown cell lines using lentivirus-based expression of specific shRNAs (Santa Cruz, Inc). We used commercial shRNAs targeting dynein heavy chain (DHC), dynactin 2 (DCTN2), dynactin 3 (DCTN3), end binding protein 1 (EB1), and BICD2. GFP control shRNA was used to determine the efficiency of lentiviral infection for each cell line. The plasmid vector also contains the puromycin resistance gene for positive clone selection. A toxicity test for puromycin was performed for SK-N-SH cell lines and it was found that 10 μg/ml was sufficient for selecting knocked down cells in SK-N-SH cell line (Fig. 4A). A serial passage of all SKNSH cells up to 10 passages did not cause significant cellular toxicity or significant replication issues. However, we observed increasing changes in morphology and viability after 10 passages with loss of cells at approximately 20 passages. Therefore, all experiments performed in this study were limited to cell lines at early passages (<5).

**Figure 4.**
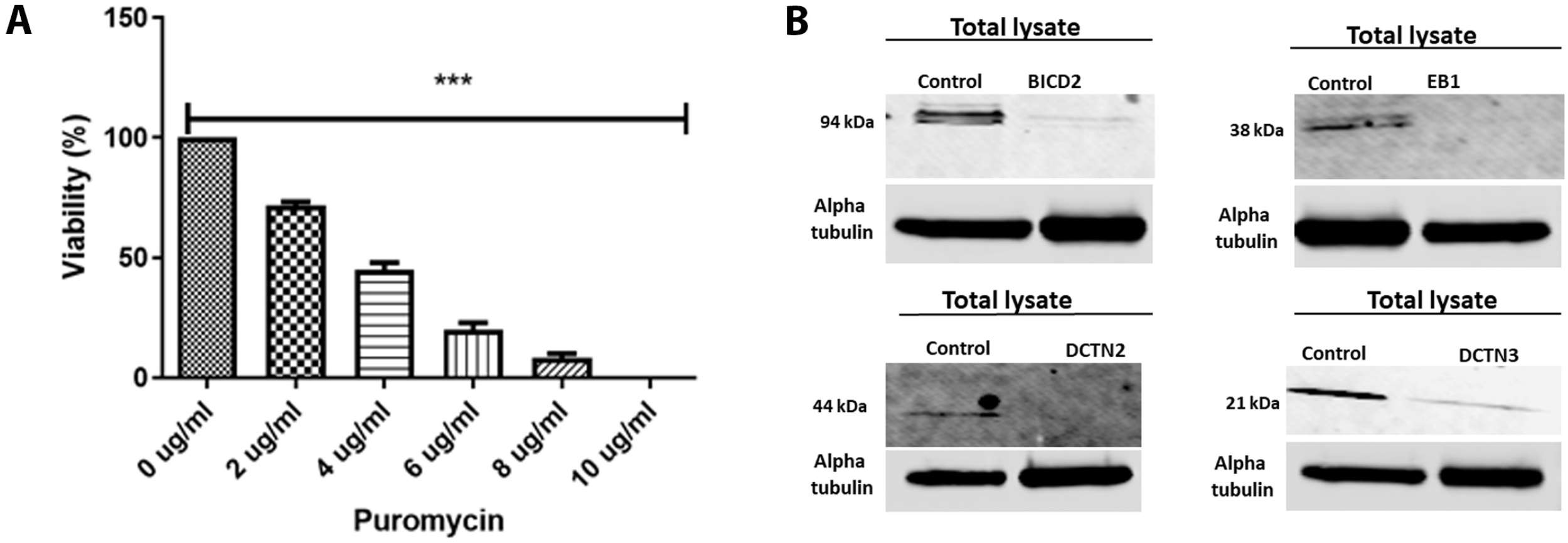
shRNA mediated silencing of Dynein cofactors in the SK-N-SH cell line. (A) Puromycin toxicity assay on SK-N-SH cells. A range of different concentrations (0ug/ml-10ug/ml) of puromycin was used to determine the appropriate amount of puromycin that is toxic to SK-N-SH cells. ***, P < 0.001 versus the no drug (0 ug/ml) treated control;statistical comparison was conducted by GraphPad prism software using ANOVA with a post hoc t-test with Bonferroni adjustment. Bars represent 95% confidence intervals about the means. (B) Lentivirus shRNA (human) mediated silencing in SK-N-SH cells. Left lanes in each panel show SK-N-SH lysate treated with control shRNA and right lanes show lysates treated with shRNA targeted against motor accessory proteins BICD2, EB1, dynactin complex (DCTN2 and DCTN3), respectively.

Cellular extracts from the engineered cell lines were prepared and the level of BICD2, EB1, DCTN2 an DCTN3 expression was assessed via western immunoblots. All cell lines showed highly significant reduction of the targeted proteins in comparison to the control shRNA treated lysates (Fig. 4B).

We investigated HSV-1 (McKrae) intracellular transport in the EB1, BICD2, DCTN2, DCTN3, and DHC knockdown SK-N-SH cells by detecting the major protein VP5 (ICP5) at early times post infection via indirect immunofluorescence assay (IFA). The SK-N-SH dynein heavy chain knockdown cells did not survive and thus were not used in these experiments. Control cells were selected using the same process as with the knockdown cell lines except that a control (scrambled) shRNA was used. Following adsorption of the virus at 4°C for an hour, control cells were incubated at 37°C for 30 min, 3 hours and 5 hours to observe the time required for the virus to travel towards nucleus. VP5 (ICP5) stained virions at 30 minutes after infection were observed predominantly in the periphery of cells. At 3 hpi, capsids were detected in the cytoplasm and proximal to the nuclear membranes. At 5 hpi, VP5 (ICP5) was detected within the nucleus representing newly expressed VP5 (ICP5) (Fig. 5A). We considered that the distinct VP5 puncta originated from intact input capsids, while diffuse VP5 fluorescence signals originated from newly synthesized VP5 that is expressed in the cytoplasm and migrates into the nuclei where virion capsids are assembled. There was a marked reduction in the major capsid protein VP5 (ICP5) detected in the nucleus in EB1, BICD2, DCTN2, and DCTN3 knockdown infected SK-N-SH cells at 5 hpi compared to the control shRNA treated SK-N-SH cells (Fig. 5B). Viral capsids were mostly localized near the plasma membrane for the DCTN2 and DCTN3 knockdown cells. In the BICD2 and EB1 knockdown cells, virion capsids appeared to be localized in the cytoplasm and nuclear periphery (Fig. 5B). Transmission electron microscopy was performed on EB1, BICD2, DCTN2 and DCTN3 knockdown infected cells. Aggregation of capsid like structures was visible in the nucleus of the control shRNA treated cells after 5 hpi. However, HSV-1 viral capsids were not observed in the nucleus of EB1 shRNA, BICD2 shRNA, DCTN2 shRNA, DCTN3 shRNA treated cells. Instead, capsids were mostly seen near the cell membrane and in the cytoplasm of infected SK-N-SH knockdown cells consistent with the IFA results (Fig. 6).

**Figure 5.**
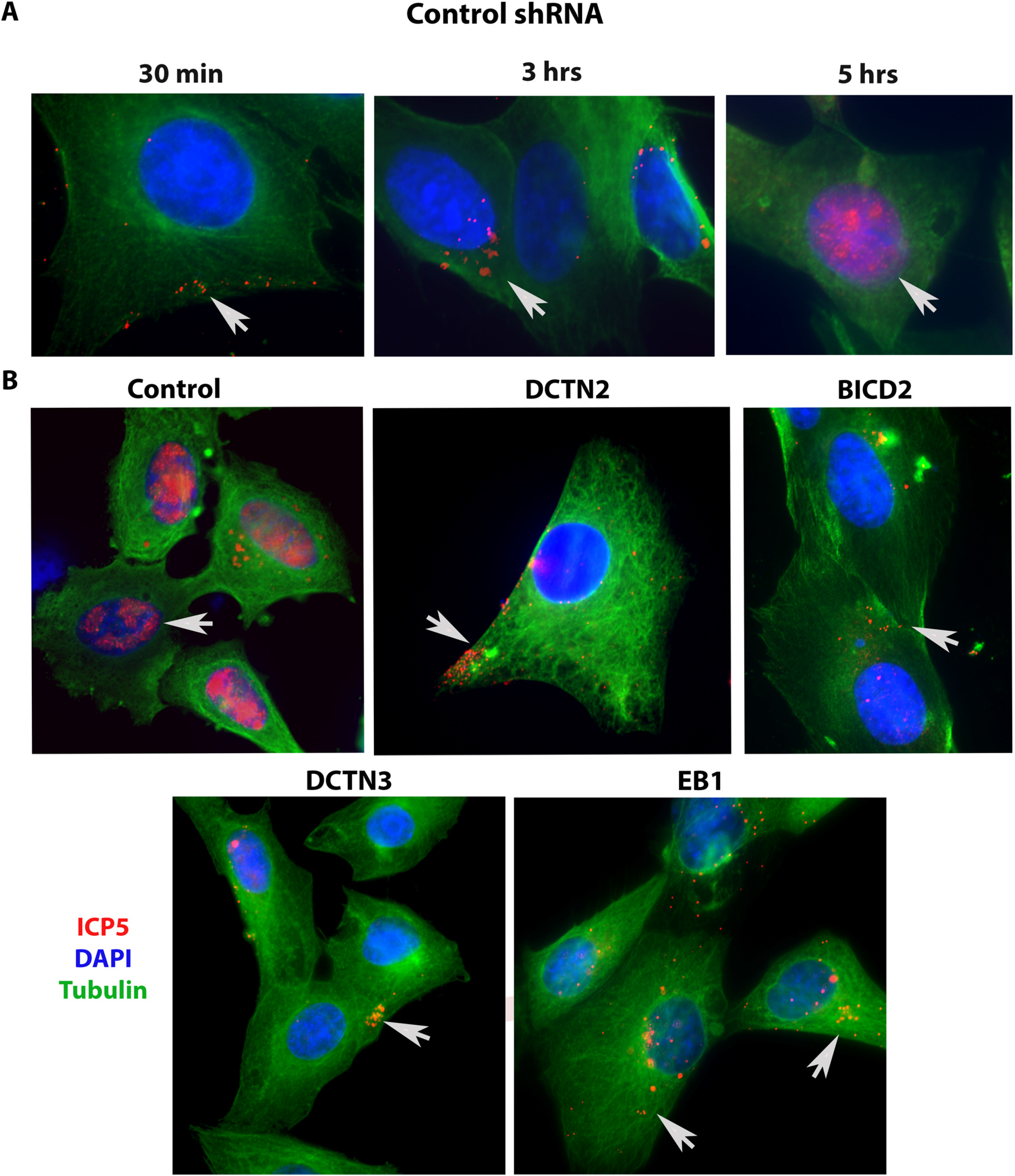
Role of motor accessory proteins in HSV-1 intracellular transport. shRNA treated SK-N-SH cells were synchronously infected with HSV-1 (McKrae) at MOI 20 for 30 minutes, 3 hours and 5 hours at 37°C. Control-shRNA treated SK-N-SH shows the presence of VP5 (ICP5) in the nucleus at 5 hpi (A). Side-by-side comparison between control shRNA and shRNA against motor accessory proteins-treated SK-N-SH cells infected with HSV-1 (McKrae) at a MOI 20 for 5 hours (B). The 8-well chamber slide was fixed with formalin, permeabilized with methanol, and prepared for fluorescent microscopy. Antibody against VP5 (ICP5) (red) was used, and nuclei were stained with DAPI (blue). Capsids colocalized in the nuclei are visible as purple (red+blue). Magnification 60X with oil immersion.

**Figure 6.**
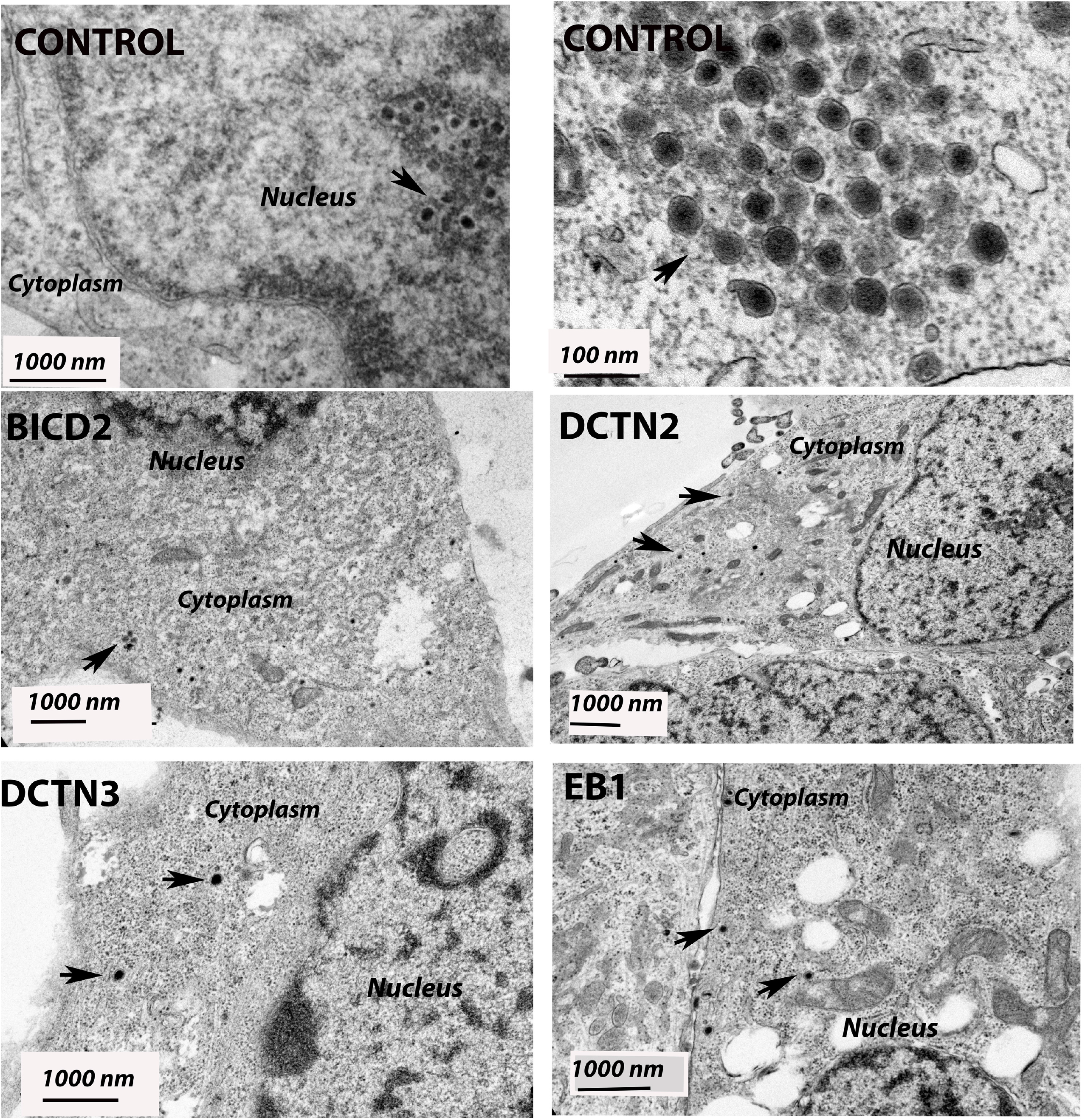
Transmission Electron micrograph (TEM) of HSV-1 (McKrae)-infected SK-N-SH cells treated with control shRNA, EB1 shRNA, BICD2 shRNA, DCTN2 shRNA and DCTN3 shRNA. The knockout SK-N-SH cells were synchronously infected with HSV-1 (McKrae) at a MOI 20 for 5 hours at 37°C. Following infection, cells were fixed and prepared for electron microscopy. SK-N-SH cells treated with control shRNA shows the presence of HSV-1 (McKrae) capsid-like structure in the nucleus. Higher magnification (100 nm) is used for better visibility. Other than control, HSV-1 (McKrae) capsids were visible only in the cytoplasm, but not in the nucleus of the motor accessory proteins knockout SK-N-SH cells. Black arrow heads specify the location of HSV-1 McKrae capsids.

### HSV-1 tegument protein UL37 and capsid protein VP5 interact with cellular motor-accessory proteins during intracellular transport

Previously, it was shown that the HSV-1 capsid interacted with EB1 and dynein and that the tegument proteins UL37/UL36 associated with dynein and kinesin motors during virion egress (30, 32, 35, 36, 38, 48, 74). To investigate whether viral tegument protein UL37 and the major capsid protein VP5 interact with the dynein intermediate chain, dynactin (DCTN1/p150, and DCTN4/p62), and the microtubule-binding protein EB1 early post infection, we performed immunoprecipitation assays. Other subunits of dynactins were not investigated due to the lack of antibodies for pull down assays. Anti-UL37 antibody precipitated DIC and anti-DIC antibody precipitated UL37 from lysates of SK-N-SH HSV-1 (McKrae)-infected cell extracts prepared at 2 hpi (Fig. 7A). Furthermore, colocalization of UL37 and DIC via immunofluorescence assay supported UL37-DIC interaction early time post infection (data not shown). Similarly, Immunoprecipitation of the same cellular extracts with anti-VP5 antibody precipitated DCTN1, p62 and EB1 but not DIC (Fig. 7B).

**Figure 7.**
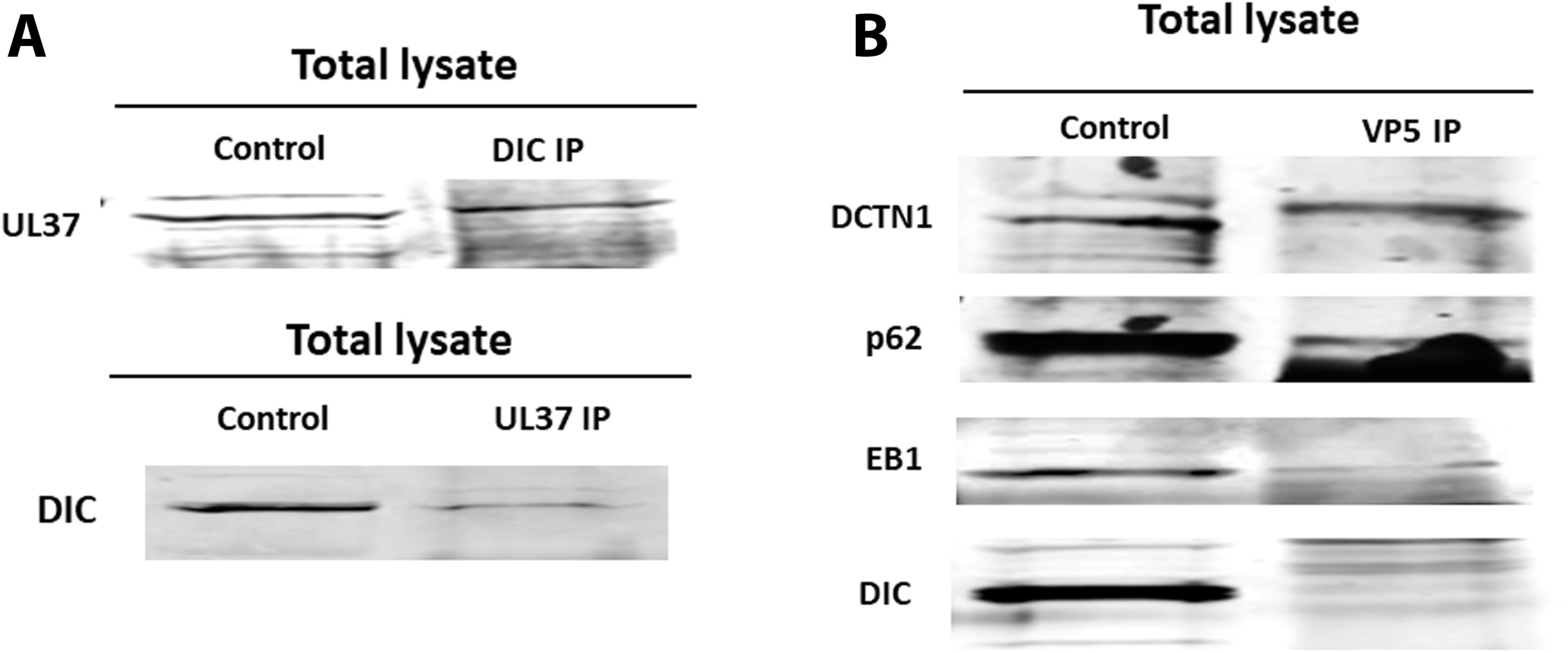
Interactions between HSV-1 capsid/tegument proteins and cellular motor and accessory proteins. (A) Two way-immunoprecipitations (IP) showing UL37 and dynein intermediate chain (DIC) interaction in HSV-1 (McKrae)-infected SK-N-SH lysates at 2 hours post infection at a MOI of 10. (B) immunoprecipitation assay showing potential interactions between VP5-DCTN1, VP5-p62, VP5-EB1 and no interaction between VP5-DIC proteins. SK-N-SH cells were infected with HSV-1 (McKrae) at a MOI of 10 for an hour and lysates were prepared with NP-40 lysis buffer with protease inhibitor cocktail. The left lane represents SK-N-SH lysates (control), and the right lane represents lysates immunoprecipitated for HSV-1 VP5 and subsequently probed for dynactin complex (DCTN1 (p150), p62 (DCTN4)), EB1 and DIC.

## Discussion

HSV-1 enters neuronal axons exclusively via fusion of the viral envelope with axonal membranes, while it can enter into epithelial cells, fibroblasts and other cells via both fusion at the plasma membrane or via endocytosis after fusion of the viral envelope with endosomal membranes (69, 75). Previously, we reported that the N terminal 38 amino acids of glycoprotein K (gK) interacts with the amino terminus of gB and modulates gB-mediated membrane fusion, and that this interaction is required for entry via fusion at the plasma membrane in neuronal axons and epithelial cells in cell culture (76–80). Furthermore, we showed that during entry into Vero and SK-N-SH cells, viral glycoprotein B (gB) interacts with Akt and triggers intracellular calcium mobilization. Interaction of gB with phosphorylated Akt, which normally resides in the inner leaflet of the plasma membrane was due to HSV induced flipping of the inner leaflet of the plasma membrane to the outer side, exposing Akt on cell surfaces (72). Lack of the amino terminus of gK prevented these events to occur and may be responsible for the inability of virions to enter via fusion at the plasma membrane. HSV-1 triggers Akt phosphorylation early times post infection in both epithelial and proliferating neuronal cell lines (65). Also, Akt phosphorylation was required for virus entry, since the Akt inhibitor miltefosine prevented virus entry (65, 72). Herein, we report that Akt and protein kinase C (PKC) inhibitors block both virus entry, and intracellular capsid transport to the nucleus in infected human neuroblastoma cells. HSV-1 infection induced phosphorylation of motor protein dynein via an Akt/PKC independent pathway, while the major capsid protein VP5 and the tegument protein UL37 interacted with the dynein motor complex and certain cofactors required for retrograde transport in SK-N-SH cells. These results show that HSV-1 entry and intracellular transport is regulated by viral protein interactions with the dynein protein complex, as wells as by modulation of kinase activities other than Akt and PKC (Fig. 8).

**Figure 8.**
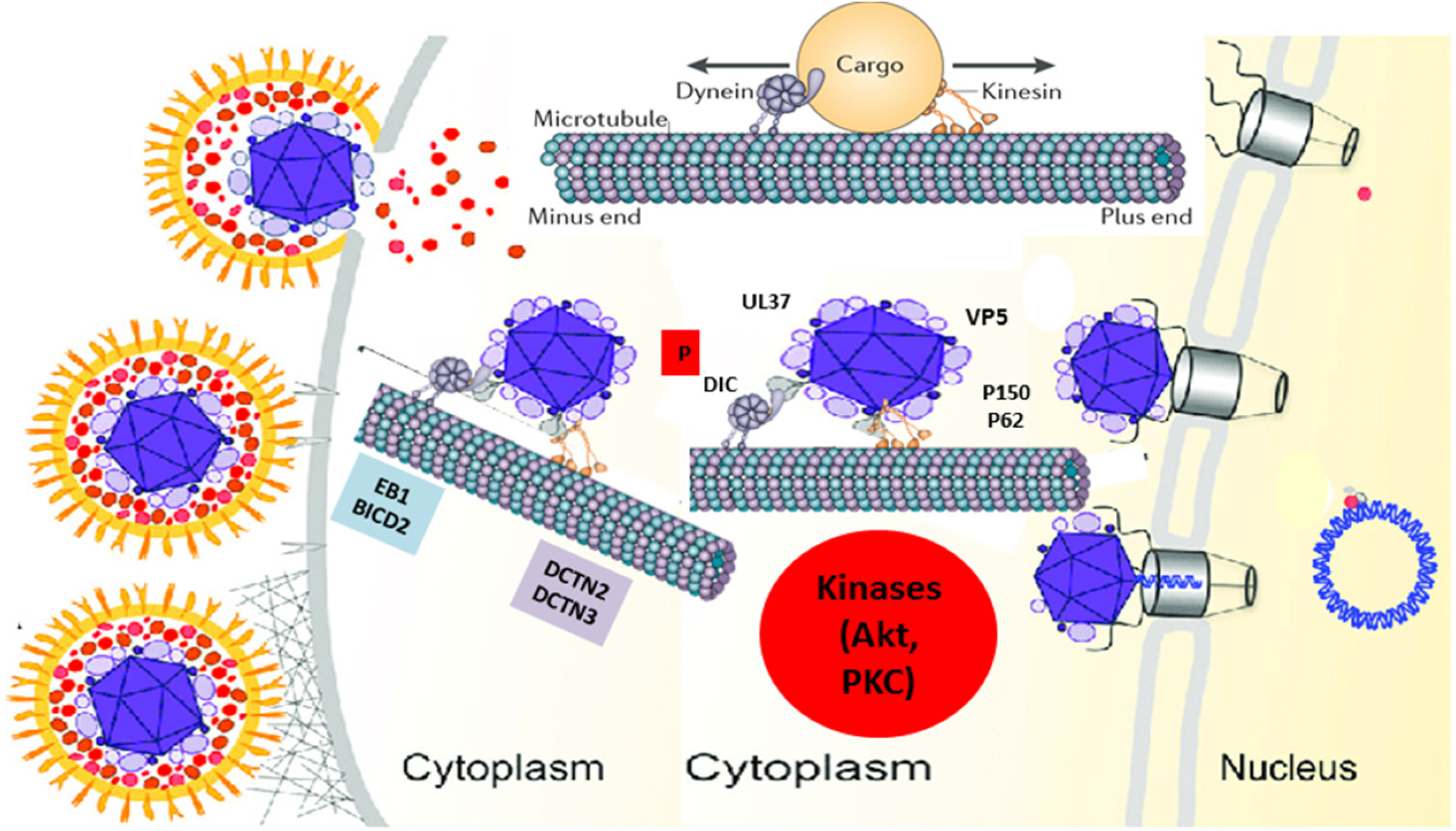
Model of HSV-1 transport into host cells. Schematic shows the molecules and intracellular signaling, which may be involved in intracellular capsid transport. Following entry into host cells, HSV-1 (McKrae) activates/phosphorylates the dynein motor and the Akt and MAPK pathways (data not shown). The major capsid protein VP5 (ICP5) and the inner tegument protein UL37 recruit the dynein motor and other accessory proteins such as BICD2, EB1, and the dynactin complex (DCTN) to facilitate retrograde transport of virion capsids. AKt and PKC activities are required for efficient entry and transport along the microtubules towards the nucleus, where the virus replicates its DNA (Adapted from Ralph *et al*., 2017, (89)).

### Intracellular signaling pathways are regulated during HSV-1 infection

Previously it was shown that HSV-1 triggers activation of MAPK pathways (ERK1/2, SAPK/JNK, p38) during late time (3 hrs, 9 hrs) post infection (66). Also, it was also reported that the HSV-1 glycoprotein H (gH) alone was sufficient to activate the JNK pathway (81). We report that activation of MAPK pathways by HSV-1 (McKrae) occurs early during infection (15 minutes post entry) (data not shown). However, virus entry or capsid intracellular transport along microtubules was independent of the MAPK pathways, since a specific MAPK inhibitor did not prevent either HSV-1 (McKrae) entry or capsid transport (data not shown). MAPK activation leads to cell proliferation, differentiation, generation of stress response, increased cell survival and motility, which may ultimately benefit HSV-1 infection (82). Also, the ERK pathway was reported to play role in the regulation of viral gene expression for SV40. adenovirus, hepatitis B virus, and HIV (83–86). Retrograde transport of NGF within signaling endosome in mammalian axons is regulated by MAPKs along with Akt/PI3K and PKC/PLC gamma (87). It will be interesting to investigate if HSV-1 induced MAPK activation indirectly regulates other kinases such as Akt or PKC.

The Akt pathway was found to be essential for HSV-1 (McKrae) entry and intracellular transport in epithelial (Vero) and neuroblastoma (SK-N-SH) cells. Importantly, our results suggest that HSV-1 (McKrae) entry and replication depend on PKC activity. The Akt inhibitor miltofesine is a potent inhibitor of Akt, however, it also inhibits kinase C. Thus, it can be concluded that kinase C is involved in both virus entry and intracellular transport of capsids. Akt directly phosphorylates GSK (glycogen synthase kinase), which regulates microtubule stabilization. Phosphorylation of different microtubule associated proteins (MAPs) by GSK3, induce accumulation of Tau, which is a characteristic of Alzheimer’s disease. GSK3 directly phosphorylates CLIP-170 but not EB1 and destabilize microtubules (88). Therefore, it is possible that Akt/PKC regulate one or more MAPs and restricts HSV-1 capsid loading onto microtubules for transport, as most of the virion capsids were seen at the periphery of the cells in SK-N-SH cells treated with an Akt inhibitor immediately following entry into cells.

### Viral activation/phosphorylation of the dynein protein complex

Following entry into host cell, virion particles can be transported either directly by recruiting cellular motor proteins, or by binding to the cargo of a specific motor protein (26). The major cellular motor protein dynein must be phosphorylated and activated to perform the motor function (9, 14–16, 23). It was shown that HSV-1 capsids require the cellular dynein motor complex for efficient capsid transport towards the nucleus (44). HSV-1 infection causes rapid dynein phosphorylation (DIC 1B S80A) following fusion/entry, which is not mediated by either Akt or PKC, since treatment with miltefosine that inhibits both enzymes did not inhibit dynein phosphorylation. Therefore, dynein must be phosphorylated by a different cellular or viral kinase. It is possible that the viral kinase encoded by the US3 gene, a structural component of the virus inner tegument, may be directly involved in dynein phosphorylation. US3 is one of three tegument proteins that remain attached to the capsid during intracellular capsids transport toward the nucleus, which places US3 in close proximity to the dynein motor machinery. Preliminary data (not shown) also suggests that US3 may interact with DIC. Future work will address whether the US3 is involved in dynein phosphorylation and the facilitation of intracellular capsid transport.

### Interaction of viral proteins with the dynein protein complex

HSV-1 utilizes dynactin and the EB1 cofactor proteins for transport towards the nucleus along microtubules following entry into host cells (7, 26, 35, 36, 43, 45). The involvement of dynactin in intracellular transport is further supported by the observation that the dynactin cofactors BICD2 is required for efficient intracellular transport as shown in HIV infection (24). Absence of dynactins DCTN2 and DCTN3, or the EB1, and BICD2 adaptors, significantly reduced HSV-1 transport following entry compared to mock (control shRNA) infection. These results suggest that HSV-1 requires dynein and its accessory proteins for efficient transport of viral capsid towards the nucleus.

In this report, we show for the first time that the major capsid protein VP5 (ICP5) directly interacts with dynactins (DCTN1/p150, DCTN4/p62) and the EB1 cofactor, which has previously been shown to facilitate HSV-1 transport by helping to load viral capsids onto microtubules (7). EB1 binds to the plus end of microtubules which is close in proximity to the cell membrane. We assume that capsid loading on microtubules via EB1 occurs immediately after entry through the plasma membrane. EB1 may detach after loading of the capsids onto microtubules or remains attached to the capsid-motor complex during the transport. Other plus end binding proteins may also be involved in this process. The tegument protein UL37 interacts with the dynein intermediate chain (DIC), which is phosphorylated after HSV-1 infection. It is not clear at this point whether DIC phosphorylation is required for interaction with the UL37 protein. Since dynactins and the dynein motor complex work synergistically in retrograde cellular transport, binding of VP5 to dynactins, and EB1, and binding of UL37 to dynein may stabilize the overall virus-motor interaction facilitating intracellular transport of capsid towards the nucleus.

Overall, these results suggest that virus entry and intracellular capsid transport require the action of cellular and viral kinases. These cellular kinases may regulate interactions of viral tegument and capsid proteins with the dynein motor machinery including dynactin and its accessory cofactors. Identification of thses these kinases and the specific protein-protein interactions that are involved may provide new ways to prevent both virus entry and intracellular transport.

## Methods and Materials

### Cells and viruses

African green monkey kidney cells (Vero, ATCC) and human neuroblastoma cells (SK-N-SH, ATCC) were used for most of the experiments. Vero cells were grown in filter sterilized Dulbecco’s modified Eagle’s media (DMEM) supplemented with 10% fetal bovine serum (FBS) and 0.2% Primocin (Invitrogen, Inc., Carlsbad, CA). SK-N-SH cells were grown in filter sterilized EMEM media supplemented with 10% FBS and 0.2% primocin. All cells were incubated at 37°C with 5% CO2. The pathogenic strain of HSV-1 (McKrae-GFP) was used throughout the experiments.

### Antibodies and reagents

Antibodies against alpha-tubulin, HSV-1 ICP5, intermediate dynein chain (DIC), dynein heavy chain (DHC), dynactin 1/p150 (DCTN 1), dynactin 2/dynamitin (DCTN 2), dynactin 3 (DCTN 3) were purchased from Abcam Inc, Cambridge, MA. Antibodies against end binding protein 1 (EB1) and VP5 were purchased from Santa Cruz Biotechnology, Dallas, TX. The HSV-1 UL37 antibody was a gift from Dr. Frank J. Jenkins, University of Pittsburgh Cancer Institute. Antibody against phosphorylated DIC was a gift from Dr. Kevin Pfister’s laboratory, University of Virginia. A MAPK family antibody sampler kit was obtained from Cell Signaling Technologies, Danvers, MA. The Akt inhibitor miltefosine and the MAPK inhibitor PD 98059 were purchased from Santa Cruz biotechnology, Dallas, TX, and the PKC inhibitors STS and Gouml were obtained from Abcam Inc, Cambridge, MA.

### Immunoprecipitation and immunoblot Assays

SK-N-SH cells were infected with HSV-1 (McKrae) at an MOI of 10. At one-hour post infection (hpi), the infected cells were lysed with NP-40 cell lysis buffer (Life Technologies, Carlsbad, CA) supplemented with protease inhibitor tablets (Sigma Inc., St Louis, MO). The samples were centrifuged at 13,000 rpm for 10 min at 4°C. The supernatants were then used for immunoprecipitation. Proteins from virus-infected cells were immunoprecipitated using protein G magnetic Dynabeads according to the manufacturer’s instructions (Invitrogen, Carlsbad, CA). Briefly, the beads were bound to their respective antibodies and placed on a nutator for 10 min, followed by the addition of cell lysates. The lysate-bead mixture was kept on the nutator for 10 min at room temperature and subsequently washed three times with 1X phosphate-buffered saline (PBS). The protein was eluted from the magnetic beads in 40 μl of elution buffer and used for immunoblot assays. Sample buffer containing five percent β-mercaptoethanol was added to the samples and heated at 55°C for 15 min. Proteins were resolved in a 4 to 20% SDS-PAGE gel and immobilized on either nitrocellulose or PVDF membranes. For western immunoblots, subconfluent SK-N-SH cells were infected with the indicated viruses at an MOI of 10 for different time points, as mentioned above. The cells were similarly lysed with the NP-40 lysis buffer, and the lysate was used for western immunoblots. Immunoblot assays were carried out using rabbit anti-Akt-1 S473 antibody (Millipore, Burlington, MA), rabbit anti-Akt 1/2/3 antibody (Millipore, Burlington, MA), rabbit anti-MAPK family antibodies (Cell Signaling Technologies, Danvers, MA), rabbit anti-dynein intermediate chain phosphorylation (DIC 1B S80A) antibody (from the laboratory of Dr. Kevin Pfister), mouse anti DIC antibody (Abcam plc, Cambridge, MA), horseradish peroxidase (HRP)-conjugated goat anti-mouse antibodies against the light chain (Fab) and heavy chain (Fc) (Abcam,plc, Cambridge, MA), and HRP-conjugated goat anti-rabbit antibody (Abcam, plc, Cambridge, MA).

### Lentivirus shRNA mediated gene silencing

shRNA targeting dynein heavy chain (DHC), BICD2, dynactin 2 (DCTN 2) dynactin 3 (DCTN 3), and end binding protein 1 (EB1) were used and delivered via lentiviral vectors in SK-N-SH cells. The level of expression of these proteins was detected by western blot analysis of lysates of transduced cells. Gene silencing by shRNA was performed according to the protocol provided by Santa Cruz biotechnology. Briefly, cells were grown in complete media (with 10% FBS and 0.2% Primocin) in 12 well plates. Lentivirus transduction was performed when cell monolayers were approximately 50% confluent. To increase the binding between the pseudo viral capsids and the cellular membrane, polybrene (sc-134220) was used at a final concentration of 5μg/ml of complete growth medium. The media was removed and replaced with 1 ml per well of polybrene media. Lentiviral particles were thawed at room temperature and mixed gently before use. The cells were transduced by adding shRNA lentiviral particles to the culture. Following overnight incubation, at 37°C, the growth medium was removed and replaced with 1 ml of complete medium without polybrene. To select stable clones expressing the shRNA, cells were split 1:3 to 1:5 depending on the cell type, and puromycin hydrochloride was used as a selection agent. Prior to transduction, a puromycin titration was done to determine the sufficient concentration to kill the non-transduced cells. 15 μg/ml puromycin for Vero cells and 10 μg/ml puromycin for SK-N-SH cells was used to kill non-transduced cells.

### Retrograde transport Assay

SK-N-SH cells were grown in eight well chamber slides and serum starved overnight. Cells were then adsorbed with HSV-1 (McKrae) at 4°C with a MOI of 20. The slides were then shifted to 37°C for 15 minutes and then washed with a low pH buffer. The cells were treated with inhibitors of choice for 5 hours at 37°C. Subsequently, the slides were fixed in formalin and permeabilized with ice-cold methanol followed by blocking with 10% goat serum. Antibody against ICP5 was used to detect viral capsid transport to the nucleus, which also was stained with DAPI. Images were taken using a 60x objective (oil immersion) on a fluorescent microscope.

### Drug toxicity/Cell viability Assay

Vero cells were grown in 12 well plates and serum starved overnight before treatment. Drugs were dissolved in DMSO, and the dilutions were made in DMEM without serum. Cells were treated with different dilutions of drugs and incubated at 37°C for 24 hours. Each dilution was tested in triplicates. The percentage of live cells was quantified using trypan blue. The counts from treated wells were compared with the non-treated wells.

### Statistical analyses

Statistical analyses were performed by using analysis of variance (ANOVA); students t-test and P values of <0.05 were considered significant. Bonferroni adjustments were applied for multiple comparisons between control and treatment groups. All analyses were performed using GraphPad Prism (version 5) software (GraphPad Software, San Diego, CA, USA).

## Acknowledgements

The work was supported by the LSU Division of Biotechnology & Molecular Medicine, by a Governor’s Biotechnology Initiative grant (to K.G.K.), and by Cores of the Center for Experimental Infectious Disease Research (CEIDR), funded by NIH, NIGMS, and grant P30GM110670. We thank Dr. Kevin Pfister for providing the phospho-dynein antibody, Dr. Xiaochu Wu for assistance with electron microscopy and Peter Mottram for helping with fluorescent microscopy.

